# Cooperative Communication with Humans Evolved to Emerge Early in Dogs

**DOI:** 10.1101/2021.01.12.425620

**Authors:** Hannah Salomons, Kyle Smith, Megan Callahan-Beckel, Margaret Callahan, Kerinne Levy, Brenda S. Kennedy, Emily Bray, Gitanjali E. Gnanadesikan, Daniel J. Horschler, Margaret Gruen, Jingzhi Tan, Philip White, Evan MacLean, Brian Hare

**Affiliations:** Department of Evolutionary Anthropology, Duke University; Department of Anthropology, Pennsylvania State University; Wildlife Science Center, Stacy, Minnesota; Canine Companions for Independence, Santa Rosa, California; School of Anthropology, University of Arizona; Cognitive Science Program, University of Arizona; Department of Clinical Sciences, College of Veterinary Medicine, North Carolina State University; Department of Psychology, University of Leiden; Department of Statistics, Brigham Young University; College of Veterinary Medicine, University of Arizona; Department of Psychology, University of Arizona; Center for Cognitive Neuroscience, Duke University; Psychology and Neuroscience, Duke University

## Abstract

While we know that dogs evolved from wolves through a process of domestication, it remains unclear how this process may have affected dog cognitive development. Here we tested dog (N=44) and wolf (N=37) puppies, 5-18 weeks old, on a battery of temperament and cognition tasks. Dog puppies were more attracted to humans, read human gestures more skillfully and made more eye contact with humans than wolf puppies. The two species were similarly attracted to objects and performed similarly on nonsocial measures of memory and inhibitory control. These results demonstrate the role of domestication in enhancing the cooperative communication skills of dogs through selection on attraction to humans, which altered developmental pathways.

**One Sentence Summary**

Domestication altered dogs’ developmental pathways to enhance their cooperative communication with humans.

## Main Text

Domestic dogs (*Canis familiaris*) rely on communicative gestures when cooperating with humans (*1, 2*), and dogs with more skill in comprehending human gestures are more successful as detection and assistance dogs (*3*). This interspecific communication is unusual: dogs are more skilled at using human gestures than mother-reared chimpanzees and other great apes (*4*–*6*). Like human children, but unlike mother-reared great apes, dogs can spontaneously use novel and arbitrary gestures (*7*–*11*). Controls reveal this flexibility is not simply explained by the use of olfactory cues, an attraction to human hands or bodily motion created by a gesture (*1, 7, 8, 10, 12*–*15*). Instead, analysis of individual differences suggests the communicative flexibility of dogs is human-like. Dogs and human infants show the same correlated variance in their use of different human gestures - a pattern not observed in other great apes (*6*).

### The Domestication Hypothesis

*(DH)* posits that the ability of dogs to understand human gestures (without intensive training) is a product of domestication (*4*). A number of lines of evidence support this hypothesis.

First, performance with human gestures in dogs varies independently from success in other cognitive tasks (*6, 10, 16, 17*) and breed differences in these skills are predicted by genetic similarity among breeds and associated with genes expressed in the brain (*17, 18*).

Second, the interspecific communicative abilities of dogs emerge early (*19*). While dogs can become more skillful using human gestures with age and training (*20, 21*), the use of human gestures does not require intensive exposure to humans. Around the age of weaning (∼7-9 weeks) dog puppies already use human gestures (*4, 8, 22*), with free ranging dog puppies and litter-reared assistance dog puppies succeeding with multiple gestures on their very first experimental trial (*11, 23*).

Third, experimental foxes selected for friendliness toward humans exhibit dog-like skills at reading human gestures. Experimental fox kits use human gestures at the level of dog puppies. They also use human gestures more than age-matched control foxes (bred irrespectively to their response to humans). They show more skill with two different communicative tasks even though the control foxes were raised with intensive exposure to humans and outperformed the experimental foxes on a non-social task (*24*). This work led to the proposal that a similar process occurred during dog domestication and led to the expression of dogs’ unusual social skills (*19, 25*).

### The Canid Ancestry Hypothesis

*(CAH)* provides an alternative to the DH and suggests instead that dogs inherited their interspecific communicative abilities from their ancestor with wolves (*4, 7, 26*). The ability of some adult wolves to learn the use of human social gestures can be viewed as support for the CAH (*22, 27*–*31*); but to date, there is limited evidence that adult wolves, even if hand raised by humans from the first days of life, show spontaneous use of human gestures as seen in dogs(*22, 27, 30*–*32*). Any skill they demonstrate likely requires intensive exposure to humans or explicit training not required for the appearance of these same skills in dogs. However, initial comparative developmental studies with dogs and wolves have yielded conflicting findings. While one comparison found that dogs but not wolf puppies spontaneously read human gestures (*22*), another found the two species performed similarly (*27*). This may suggest the early emerging skill of dogs is inherited from a common ancestor with wolves or that the second comparison was not sensitive enough to detect a significant developmental difference between the species (i.e. this experiment only included a small sample of wolf puppies, N=6, because the rest were too aggressive to test, Gácsi et al., 2009).

What is urgently needed is a large-scale comparison of wolf and dog puppies on a battery of cognitive tasks that includes social (especially those requiring the use of human gestures) and non-social problems. The DH predicts that dog puppies, regardless of age or amount of human interaction, will be attracted to humans more than wolves and will outperform wolves using the two human gestures but not in the nonsocial tasks. In contrast, the CAH predicts the amount of exposure to humans will be related to performance on social cognitive tasks in both species (i.e. older dogs should outperform younger dogs and human raised wolves should outperform the youngest dog puppies still living with their mother and littermates).

Here we provide this critical test by comparing the temperament and cognition of the largest sample to date of dog and wolf puppies between 5-18 weeks old (Duke University IACUC #A105-17-04). All dog puppies were retrievers bred and raised for assistance work. Most (94%) wolf puppies were only the first (56%) or second (37%) generation bred in captivity. Wolf puppies remained with littermates but received 12-hour (24%) or 24-hour (76%) human care from 10-11 days after birth. This included caregivers remaining available for constant contact, feeding the puppies by hand and sleeping with them each night up to and throughout the testing period. One of the experimenters who helped raise the wolves was always present during testing. In comparison, the dog puppies received far less contact with humans. All dog puppies remained with their mothers until weaning around 6 weeks of age, and with their littermates until ∼8 weeks of age. During this time, they mainly interacted with humans during short routine caretaking tasks. Around eight weeks of age, puppies were then sent to live with human families. The majority of dog puppies (N=27) were tested at 7-8 weeks of age, prior to going to live with human families, while a subset of puppies (N=18) were tested between 10-17 weeks of age, while living with a human family. As part of their training, dog puppies did not sleep with humans at night.

We first ran a temperament test in which subjects could approach an unfamiliar or familiar human or object to retrieve food. The unfamiliar human was an experimenter whom the puppy had never met, while the familiar human had calmly interacted with the puppy for at least 30 minutes. The unfamiliar object was a novel toy (a plastic bear) and the familiar object was taken from a subject’s enclosure (e.g. a plastic bottle, etc.). We measured how often subjects touched each stimulus in two sessions of four 4-trial blocks yielding a total of 32 trials. Using a linear contrast test on a mixed-effects logistic regression model reveals that relative to a wolf puppy, the dog puppies’ odds of touching the unfamiliar human, familiar human, and unfamiliar object were 30.52 (95% CI 14.48-73.68), 5.36 (95% CI 2.99-10.06), and 1.91 (95% CI 1.16-3.20) times higher respectively (**Fig. 1**).

**Fig. 1.**
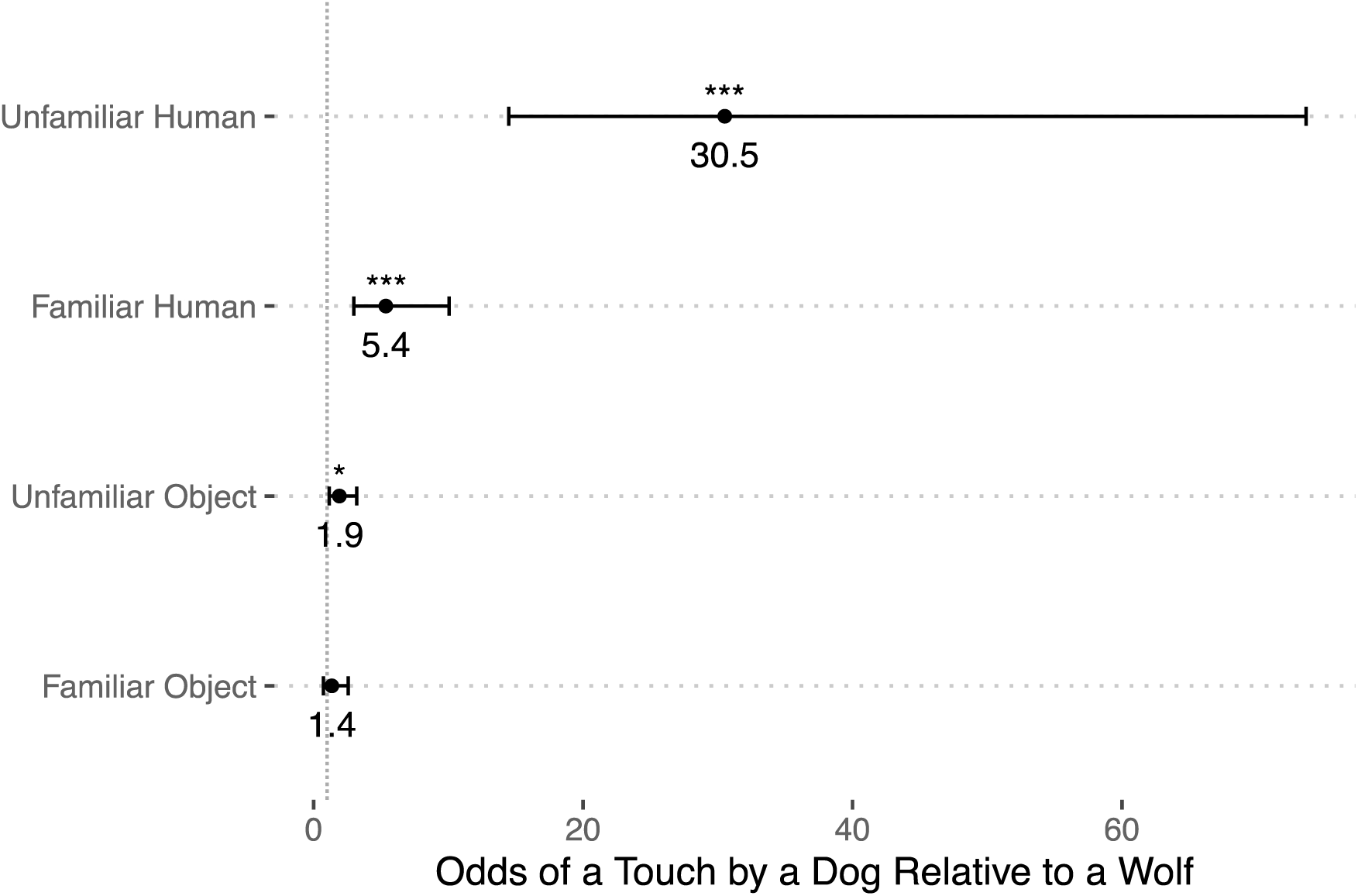
Temperament test - odds that a dog would approach and touch the stimuli in each condition as compared to a wolf (i.e., a dog’s odds of touching the unfamiliar human are 30.5 times higher than those of a wolf). *** indicates p<.001, * indicates p<.05. Vertical dotted line (odds =1) signifies the point of no difference between species. Bars signify the 95% confidence interval.

We then tested subjects’ memory for where they had recently seen food hidden. Subjects watched from 2m away as an experimenter hid food in one of two bowls separated by 2m. Once the food was hidden, the experimenter, sitting centered between the bowls, lowered her head and remained motionless while the subject was released to search for the food treat. A choice was scored when the subject approached either bowl closely enough that the nose passed over the bowl’s edge. We measured how many trials it took each subject to correctly choose the baited bowl in four out of five consecutive trials. On average, the two species took approximately the same numbers of trials to meet this criterion (**Fig. 2a**).

**Fig. 2.**
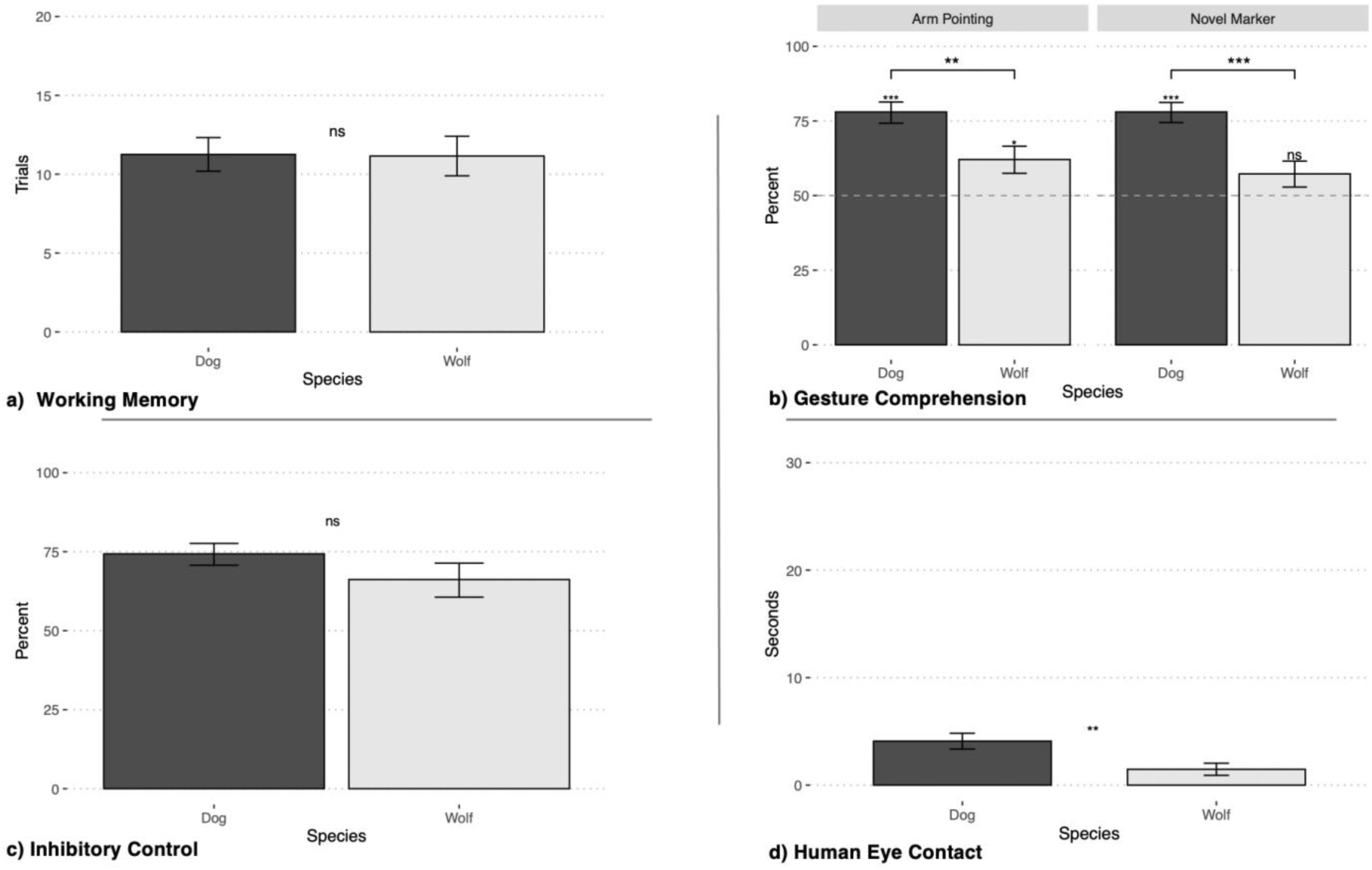
Cognitive tasks - A) Mean number of trials (±SEM) subjects required to reach criterion; B) Estimated probability (±SE) of subjects choosing the location indicated by the experimenter’s gesture, with dashed line at the level of chance (50%); C) Estimated probability (±SE) of subjects retrieving the food by navigating directly to the side opening without first touching the clear cylinder; D) Mean number of seconds (±SEM) subjects spent looking at human face during 30 second trial. For all panels, * indicates p<.05, ** indicates p<.01, *** indicates p<.001.

Subjects were then tested for their understanding of human gestures using the same method as the memory test, except the experimenter sham baited the bowls so the subject knew food was hidden but did not know where, and then gave a communicative gesture indicating the food’s location before releasing the puppy. The experimenter gestured by sitting centered between the bowls and either 1) pointing with an extended arm while gazing at the food location or 2) placing a physical marker (small ordinary wooden block) next to the correct location. Subjects received 6 trials with each gesture as well as 6 trials with each of the two control conditions. The controls tested 1) subject’s preference between two competing social cues: the location of a human near one bowl and a pointing gesture toward the other bowl, and 2) their ability to find hidden food with only olfactory information. Using a mixed-effects logistic regression model with random intercepts for each subject, we estimated the probability of choosing the hiding location an experimenter gestured towards for each species using linear contrasts. The dog puppies were estimated to choose the indicated location 77.99% of the time (95% CI 70.26 – 84.17, z = 6.11) for the pointing gesture and 78.01% of the time (95% CI 70.73 – 83.89, z = 6.46) for the marker gesture, while wolf puppies were estimated to choose the indicated location 62.08% of the time (95% CI 52.86 – 70.51, z = 2.55) for the pointing gesture and 57.25% of the time (95% CI 48.59 – 65.50, z = 1.64) for the marker gesture. The dog puppies as a group performed above chance (50%) for both the pointing gesture (p<.001) and the marker gesture (p<.001) but not the controls. The wolf puppies as a group performed above chance for the pointing gesture (p =.011) but not the marker gesture (p=.100) nor the controls. For both gestures, the dog puppies chose the indicated location significantly more than the wolf puppies did (pointing p=.006, marker p<.001) **(Fig. 2b)**. Using a linear contrast test on a binomial logistic regression model reveals that relative to a wolf puppy, a dog puppy’s odds of choosing the indicated location for the pointing gesture and the marker gesture were 2.08 (95% CI 1.14 – 3.77) and 2.58 (95% CI 1.42 – 4.68) times higher respectively (**Fig. S3)**.

As individuals, 17/31 dog puppies and 0/26 wolf puppies performed above chance when combining their performance with both gestures [≥10/12 correct using a significance threshold of p<.05, binomial probability]. On their very first trial dog puppies used both gestures [pointing and marker: 28 of 31 correct, p<.001, binomial test] while wolf puppies were not above chance on their first trial [pointing: 18 of 26, p = .07, marker: 15 of 26, p=.56]. Using a binomial logistic regression, we modeled the effect of trial number on performance and found no evidence of learning within the test in either species [pointing p=.15, marker p=.28; see **Tables S7-S8**]. In the two control conditions both species performed at chance levels ruling out either an aversion to approaching a hiding location near a human or a reliance on olfactory information. Using linear regression models including age and sex as covariates, species was the most significant predictor variable for performance on both gesture tests, and including the age parameter did not improve either gesture model for either species (**Tables S1-S2**). Finally, we found that number of approaches in the temperament test was positively associated with performance in the pointing and marker gesture comprehension tasks but not the controls. However, the relationship between temperament and gesture comprehension did not hold once species was included in the model, suggesting that the relationship was due to the species difference in temperament and not individual variation (**Tables S9-10**).

We measured subjects’ inhibitory control using a task in which an experimenter placed food inside a transparent cylinder as subjects watched. Subjects were released to retrieve the food by reaching their muzzle into one of the open ends of the cylinder. Because the cylinder was oriented perpendicularly to them, subjects needed to walk around to the open side to access the food. First, they were familiarized with this navigation on an opaque cylinder; then the cover was removed and they still needed to walk around to the side opening despite now being able to see the food directly in front of them through the transparent cylinder wall. In ten test trials, subjects’ responses were coded as correct when food was obtained by making a clear route around to one of the open ends, or incorrect when the subject first attempted to obtain the food by touching the transparent wall of the cylinder. The two species did not differ in their performance in the inhibition task (**Fig. 2c**).

As a secondary social measure, a subset of subjects (Dog: N=34; Wolf: N=15) were assessed for their propensity to make eye contact with a human experimenter. Subjects watched as food was placed into a container which they had previously opened and successfully obtained food from; however, it was then secured so they no longer could open it. In 4 trials of 30 seconds each we measured the amount of time subjects spent looking at the experimenter after encountering the box they could no longer open on their own. Dog puppies made significantly more eye contact with the experimenter than wolf puppies [Dog: M±SD =4.09±4.29 seconds; Wolf: M±SD =1.47±2.18 seconds, t(45.9) = 2.83, p = .007, Welch independent t-test] and a linear model shows that species is a significant predictor for duration of eye contact (**Fig. 2d**).

Sex, age and trial number were considered as covariates for each of the temperament and cognitive tests using linear regression model comparison, and species was always the most highly significant predictor variable on tasks where a significant species difference was found (**Tables S1-S8)**. Including age as a covariate did improve the models for all tests except the gesture comprehension tests (arm pointing and marker), further reinforcing that this ability is early-emerging in puppies (**Tables S1-S6**).

Results support the predictions of the Domestication Hypothesis. Dog but not wolf puppies are attracted to humans and show early emerging skills for reading human gestures, even though the wolf puppies received more intense human socialization. Dog puppies were more than twice as likely than the wolf puppies to use both human gestures correctly. Half of the dog puppies were successful at the individual level while no wolf puppy was. Dog puppies also spontaneously used both gestures on their first test trial and there was no evidence for increased success within test sessions or in older puppies. The youngest dogs also spontaneously used cooperative-communicative gestures despite having far less human exposure than the wolf puppies.

Success with the arbitrary marker rules out that dogs only use gestures due to an attraction to human hands, because in this condition the human placed their hand next to both of the possible hiding locations. The chance-level performance of wolves on the marker gesture rules out an aversion to the block influencing their choices. Chance performance in the body position control rules out an aversion of approaching humans explaining the wolf puppy’s performance. Wolves did not avoid the location adjacent to the pointing experimenter. The olfactory control rules out the possibility that subjects found the food rewards using olfactory cues.

Performance on the non-social tasks also suggests that relative to wolves, dog puppies are specialized for cooperative communication with humans (*6, 33, 34*). Both species performed similarly on the memory and inhibition tasks, suggesting that species differences in early ontogeny are predominantly in the social domain (with any species differences in these non-social skills likely appearing later in development, Marshall-Pescini, Virányi, & Range, 2015). Our supplemental social task further suggests that dog puppies are prepared to communicate with humans since they made more eye contact than the wolf puppies, though further development may then interact with species differences in persistence or enhance the use of human gaze (*36, 37*). And lastly, results from the temperament tests replicate similar findings (*38*–*41*) and support the idea that it is an unusual interest in humans that motivates the early emerging social skills of dogs. Dogs’ odds were 30 times higher than wolves’ to approach a stranger and 5 times higher to approach a familiar caretaker. This approach behavior was linked to proficiency using human gestures in a similar way as has previously been observed in experimentally domesticated foxes (*24*). Together, these results suggest that as wolves came under selection for friendlier behavior toward humans, developmental pathways were targeted so that the first proto-dogs began cooperatively communicating with humans in a new way. This initial evolution may have led to a second wave of selection directly acting on individual differences in cooperative communication (*42*).

## Acknowledgements

The authors wish to the thank the staff and volunteers at the Wildlife Science Center and Canine Companions for Independence for making this research possible. We also wish to thank Kerri Rodriguez, Leah Kaiser, Ben Allen, Kylie Gradie, James Brooks, Christina Burt, Charlotte Burnham, Anya Parks, and Sam Lee for their help with data collection.

## Data and materials availability

All data, code, and video used in the analysis is available upon request to any researcher for purposes of reproducing or extending the analysis.

