## Supplementary Materials for "Cooperative Communication with Humans Evolved to Emerge Early in Dogs"

**This PDF file includes:**

Materials and Methods
Supplementary Results
Figs. S1 to S3
Tables S1 to S11

### Materials and Methods

Subjects

A total of 44 dog puppies and 37 wolf puppies participated in cognitive and temperament testing. See Table S11 for the species, sex, population, testing year, rearing experience, and age at each test for each subject. All dog puppies came from lines of dogs being bred and socialized for work as assistance dogs (see MacLean & Hare, 2018 for details). Assistance dogs are not particularly skilled at using human gestures relative to other populations of dogs. MacLean and Hare (2018) found that a large sample of adult dogs from Canine Companion for Independence were less skilled as adults in their use of a human pointing gesture in comparison to a heterogenous population of pet dogs and military working dogs (see Fig. S2). Although this made it less likely we might see a difference between the dog and wolf puppies we compared, we decided to test assistance dogs as our comparison group because they offer ease of access to large numbers of same-aged puppies, and relatively high quality information available regarding their breeding and rearing histories. This means all dogs were Labradors, golden retrievers or Labrador golden crosses with known pedigrees.

The majority of dog puppies were tested at Canine Companion for Independence (CCI; www.cci.org) Headquarters in Santa Rosa, CA. The other puppies were tested at the Duke Canine Cognition Center (DCCC) in Durham, NC. The puppies tested at CCI Headquarters (N=27) were all born in Northern California, either at the Canine Early Development Center on CCI's campus (N=6) or in the homes of volunteer breeder caretakers (N=21). Puppies stayed in a whelping pool with their mother for the first three weeks, and then moved to a larger pen with their mothers until they weaned at ~6 weeks of age. During this time, puppies had relatively limited interaction with human caretakers (<2 hours total per day), who mainly moved pups while cleaning the pool and pen, cut toenails, took daily weights, and provided food and water for the mother. After weaning, puppies continued to live and sleep in a large floor pen with littermates, either in the whelping center or inside a volunteer home, and humans provided kibble three times per day (see Bray et al., 2020 for details). While puppies had multiple hours of exposure to humans each day, their living set-up (i.e. floor pens) allowed for less prolonged interaction with humans as compared to the wolf puppies. Furthermore, unlike the wolf puppies, dog puppies only ever slept with conspecifics.

Additionally, a few CCI puppies were tested at the DCCC (N=5). These puppies had the same breeding pedigree and rearing experience as those tested at CCI Headquarters, and were then sent to live at Duke Puppy Kindergarten (DPK) when they were ~10 weeks old. At the DPK, the puppies lived together and spent their days in an indoor playroom, outdoor playpen, and on brief outings around campus. They had human caregivers with them all day and were occasionally visited by adult CCI dogs for play sessions. They slept separately in kennels without humans at night.

Finally, a minority of dogs were recruited from two local assistance dog nonprofits that breed and raise dogs near Duke University: Paws 4 People (P4P; www.paws4people.org; N=8), and Ears Eyes Nose & Paws (EENP; www.eenp.org; N=5). Similarly to the CCI puppies, these dogs were raised in human homes with their mothers and litter mates until ~8 weeks. They were then weaned and reared with their littermates in the homes of the staff until our testing was completed. These staff raisers brought the puppies to DCCC for testing.

Dog puppies were selected for testing solely based on their age and availability. We attempted to test 44 dog puppies and were able to test all 44 (although we aborted testing after the first temperament session for one puppy, Idris, due to low motivation).

All wolves were born and raised at the Wildlife Science Center in Stacy, Minnesota (WSC; www.wildlifesciencecenter.org). All the wolf puppies tested were either the first (N=21), second (N=14), or third (N=2) generation to be born in captivity, and known to have wild North American ancestry. The wolves were selected for testing based on their availability, age and willingness to participate in tasks involving human interactions. The wolves were taken off their mothers at 10-11 days of age (not for the purpose of this study) and raised by humans, alongside their littermates and adult dogs. 28 wolf pups had human and adult dog contact 24 hours per day 7 days a week until all testing was completed: at least one human raiser and at least one adult dog were always inside their enclosure, even at night time, when the pups often slept next to or on top of the human raiser. The other 9 wolf puppies were in constant human and adult dog contact during the day (11-12hrs per day), but at night were only with other same age wolf pups, until testing was complete. All wolf pups were bottle fed by their human raisers until at least 8 weeks of age, with some raw ground meat introduced by hand during this period as a supplement. After 8 weeks, they continued to eat raw meat (fresh ground and/or carcasses) provided by their human caretakers and were occasionally offered bottles for comfort as needed.

We attempted to test a total of 49 wolf puppies and were able to collect data from 37. We were unable to collect any data with the other 12 wolf puppies because they were too nervous around unfamiliar humans (i.e. experimenters), even though they had been heavily exposed to humans: all 12 were hand raised after 10-11 days with their mothers, with 3 being raised with approximately 12 hours per day of human and dog contact, and 9 being raised with approximately 24-hour contact. There was always a human raiser and a familiar adult dog present during testing, but even so, they would hide, pace or refuse to eat when an unfamiliar human was in the testing room. Testing of these 12 subjects was abandoned after failing in multiple attempts to habituate them to the presence of a novel experimenter, as they were unwilling to approach the testing area, make choices, or search for food.

Not all subjects participated in all tests, largely due to time constraints and logistics of collecting data over multiple years at two field sites located in different regions of the United States. This required flights and defined data collection periods that matched the required stages of development being tested. This was particularly difficult in the case of the wolves, which have one whelping period in spring as opposed to dog puppies, which are born year-round. We completed the entire battery of five tests (i.e. temperament, working memory, inhibition, gesture comprehension, and human eye contact test) with 8 wolf pups and 28 dog puppies. Nine additional wolf pups completed all tests except the human eye contact test (which we only began implementing with the wolves in 2017).

In both species we were able to balance for the number of males and females tested. We also made sure that the population of wolves we tested were not younger than the dogs tested. This ruled out explaining any performance by the dog puppies that was significantly better than the wolves as a result of the dogs being more mature developmentally. For all tests, the mean age of the wolves was older than that of the dogs. We also sampled from a wider range of ages in the dogs than the wolves, with the one exception being in the inhibition test where two 5-week-old wolves were included.

This research was approved by the Duke University IACUC #A105-17-04.

Setup of Testing Area

Dog puppies were tested in a quiet testing room, at either CCI or DCCC as described above. Before the first testing session, all the pups tested at CCI were unfamiliar with the room. All EENP and P4P pups tested at DCCC were also unfamiliar with the room prior to testing. The 5 CCI puppies raised in DPK and tested at DCCC were familiar with the room before testing. All wolves were tested in a familiar room at WSC, adjacent to their outdoor living area, in which they had spent significant time. While background environmental sounds (such as dogs barking or facility trucks driving on gravel) could be heard from within the room, these were sounds that the wolf pups heard all day throughout their lives and rarely reacted to during testing; if the wolf puppies seemed distracted by ambient noise, a pause was taken between trials. All dog and wolf puppies, regardless of prior familiarity with the room, were given a few minutes to explore and acclimate to the room before testing began. While testing room familiarity was different between species, dogs have performed similarly on these types of tests in both familiar and unfamiliar test spaces (*10*). The unfamiliarity of the testing spaces for most dog puppies is conservative since it potentially works against the experimental hypothesis (i.e. if anything, dog puppies might perform less skillfully in new surroundings).

The standardized testing area (**Fig. S1**) was delineated using chalk (WSC), tape (CCI HQ), or testing mats (DCCC). The setup for this study consisted of two 2 m lines on the floor arranged in the shape of a capital “T”, which intersect at position C. Positions L and R were 1m to the right and left of position C, and positions R’ and L’ were 20cm to the right and left of position C. At CCI HQ, the testing area was fenced in within the larger room by a puppy pen. At WSC and DCCC, the room was partitioned into a smaller area by use of a temporary wall (DCCC) or wire fencing (WSC). All testing sessions were recorded by two or three handheld video cameras (Sony Handycam HDR-CX405 or similar), mounted on tripods and positioned outside the testing area at locations which best captured the behaviors of the subject and experimenter for each test.

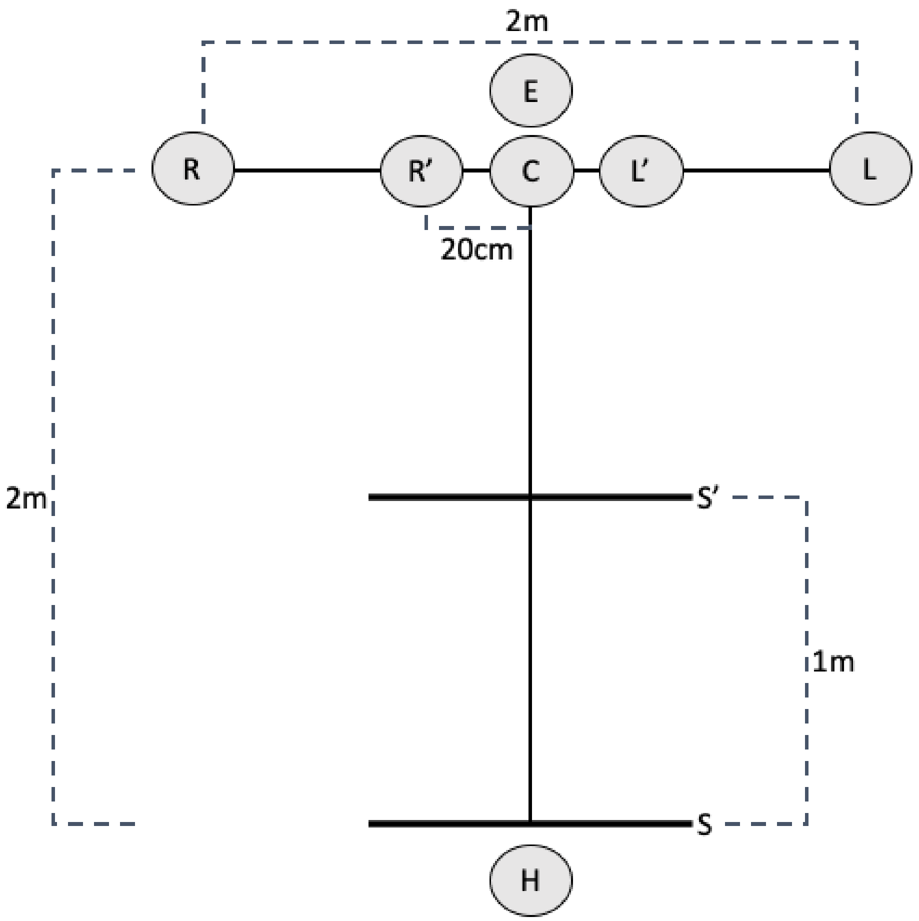


**Fig. S1 Testing Area**

**<u>Temperament Test</u>**

<u>General Procedures:</u>

Subjects were presented with the opportunity to retrieve a piece of food nearby a familiar or unfamiliar human or object. For the dog puppies, the familiar human was someone to whom the subject had at least 30 minutes of prior exposure. For the puppies tested at CCI, the person who would play the "familiar human" role became familiar with the subjects on the morning of their testing day. The familiar human entered the pen where the subjects lived with their littermates and engaged in whatever way the subjects showed interest (sitting with and talking to, petting, playing, etc.) for 30 minutes. For the puppies tested at DCCC, the familiar person was one of their regular caretakers. For the wolf pups, the familiar human was always a hand-raiser. For both species, the unfamiliar human was a stranger the subject had never seen before. The familiar object for both species was a toy that the subject had prior exposure to and had been observed interacting with in their living area, while the unfamiliar object was a remote-controlled plastic bear the subjects had never seen before. Each stimulus was presented in four 4-trial blocks for a total of 16 trials during each session, counterbalanced for order across individuals in four different session orders (familiar/unfamiliar first, and human/object first). The experimenter (E) presented the food and the stimuli, and the handler (H) positioned the subject and released the subject at the appropriate times during trials. Prior to the stimuli presentation, E always baited the platform (overturned bowl) with a piece of food at location C. In each four-trial block, the stimuli were always placed at positions R, L, R' and L' (**Fig. S1**) in that order.

*Warm-up Trials:*

Warm up trials were done prior to completing any test trials to introduce the set-up to the subject. H centered the subject at the start line (S) and gently held the subject in place. E approached the subject to present the food reward in her hand and said "look!", allowing the puppy to see and sniff the food briefly. E walked backwards (facing the puppy with food visible in hand) to set the reward on top of the platform at location C, then exited the testing area (or walked to the back of the room if the testing area was not fenced) and stood still and silent, facing the wall. Once E was positioned, she said "okay!", signaling H to let go of the subject. The subject had 30 seconds to approach the platform and retrieve the food reward. If the subject did not approach the platform within 30s, the trial was repeated. The subject was required to successfully retrieve the reward on two consecutive trials in order to advance to testing. If the subject did not show any interest in approaching the food at first, a warm-up trial was done with the food placed directly in front of the subject, then halfway between the subject and position C, and then the standard warm up trials were completed at position C.

Session 1: Stationary Stimuli

*Familiar and Unfamiliar Object Trials:*

Object trials were conducted in the same way as the warm-up trials, with the addition of the object presentation. The trial began as described for the warm-ups, but after E placed the food on the platform, E then retrieved the appropriate object for the trial (familiar or unfamiliar) from a table within reach but outside the testing area. E approached the subject again with the object in hand, bent down and presented it a few inches from the subject's face, and said "look!". E walked backwards, facing the puppy with the object visible in hand, to place the item at the appropriate position (R, L, R' or L'). E then exited the testing area and stood still and silent, facing the wall. Once E was positioned, she said "okay!", signaling H to let go of the subject. The subject had 30 seconds to approach the platform and retrieve the food reward and/or interact with the object if desired. During the trial, H sat still in place, watched the subject without interacting. She said "Food" aloud in a neutral tone if the subject ate the food, and "Touch" if the subject touched the object with its nose, mouth, or front paws. This allowed E, who could not see the subject during the trial, to record these responses on the data sheet, and was helpful for reference in the videos (although we only used whether or not the subject touched the stimulus in the final analysis) . Regardless of the subject's behavior, the trial ended after 30 seconds, and the subject was not given the food reward if she had not retrieved it.

*Familiar and Unfamiliar Human Trials:*

The human trials were exactly the same as the object trials, except that the stimulus was a human instead. After E placed the food on the platform, E then knelt at the appropriate position (R, L, R' or L'), facing towards C, and looked down with her hands flat on her thighs. Once E was positioned, E said "okay!", signaling H to let go of the subject. E remained in position for the duration of the trial, which proceeded in the same way as the object trials described above.

Session 2: Moving Stimuli

*Warm Up Trials:*

Refamiliarization trials, identical to the warm-ups of Session 1, were conducted if there was more than a 30-minute break between Session 1 and Session 2. Most subjects completed these sessions consecutively and refamiliarization was not required.

*Familiar and Unfamiliar Object Trials:*

Session 2 trials were identical to Session 1 except that the stimuli moved as described below before the dog was released to make a choice:

Familiar Toy: After placing the object at the appropriate position (R, L, R’ or L’), E stood behind the object and, keeping her eyes looking down at the ground, pulled on a string attached to the object to lift it off the ground to about knee height, then quickly lowered it back to its place on the floor. E lifted and lowered the toy this way three times before exiting the testing area.

Unfamiliar Toy: After placing the object (a plastic bear) at the appropriate position (R, L, R’ or L’), the bear moved three times. For some of the initial subjects E stood behind the bear looking down while the bear turned 360 degrees via remote control but for most subjects E simply moved the bear up and down using a string.

*Familiar and Unfamiliar Human Trials:*

Session 2 trials were identical to Session 1, except that the familiar or unfamiliar human playing the role of E moved her upper body after kneeling at the appropriate position (R, L, R’ or L’). E bent her upper body down as much as she was able, so that her torso approached her thighs and her forehead nearly touched the ground, placing both hands on the ground underneath her shoulders, and then pushed herself back up. She performed this unusual movement a total of three times before resuming the same still position as in Session 1.

<u>Abort Criteria</u>

If the subject exhibited any signs of significant stress (including excessive whining, barking, escape behavior, or defecation), a break was taken. If time allowed, another attempt occurred later the same day. After a break, the subject repeated the warm-up trials before resuming.

**<u>Working Memory:</u>**

<u>Warm-Ups - Visible Placement:</u>

At the beginning of the session, the subject was required to pass a warm-up criterion prior to completing any test trials. Warm-up trials were conducted to assure that the subjects were motivated to search for the reward and to prevent side biases. Warm-up trials consisted of two phases: (1) one-bowl centered visible placement (2) one-bowl alternating visible placement.

*Phase 1 - One Bowl Centered:*

H first centered the subject at the start line (S) and gently held the subject in place. E approached the subject to present the food reward in her hand and said “look!”, allowing the puppy to see and sniff the food briefly. E walked backwards, facing the puppy with food visible in hand, to gently set the reward into the food bowl at location C without making a sound. After baiting, E knelt behind the bowl and rested her hands flat on her thighs, looking straight down at her lap. The subject was then allowed to approach the bowl and obtain the reward. If the subject did not approach the bowl within 15s, the trial was repeated. This phase of warm-ups familiarized the subject with the set-up and assured that the subject was motivated to find the reward. To pass the warm-up criteria the subject was required to successfully retrieve the reward from the bowl on two consecutive trials within a maximum of six trials.

*Phase 2 - One Bowl Alternating*

Phase 2 warm-ups were identical to phase 1, except that the bowl's position was counterbalanced between the R or L positions. If the subject did not approach the bowl in 15s, the trial was repeated. Repeating these trials served as a correction procedure for spontaneous side biases and ensured that subjects gained experience finding the reward in both locations. To pass the warm-up criterion the subject was required to successfully retrieve the reward from the bowl on two consecutive trials (one on each side) within a maximum of six trials.

Working Memory Test Trials:

Bowls were placed at both positions R and L, and E was provided a list denoting which side to place the food on for each trial. Side placement was counterbalanced with the constraint that the reward could not be on the same side in more than two consecutive trials. H centered the subject at the start line (S) and gently held the subject in place. E approached the subject to present the food reward in her hand and said "look!", allowing the subject to see and sniff the food briefly. E walked backwards, facing the puppy with food visible in hand, to gently set the reward into the food bowl at the appropriate location (R or L) without making a sound. After baiting, E walked back to the center (location E), knelt, and rested her hands flat on her thighs, looking straight down at her lap. The subject was then allowed to approach the bowl and obtain the reward. If the subject did not approach the bowl within 15s, the trial was repeated. If the subject chose the baited bowl, the subject was allowed to have the reward and the next trial was administered. If the subject chose the incorrect bowl, E said "wrong" in a monotone voice and the subject was not rewarded nor allowed to see where the food was located. If the subject did not choose any bowl within 15s, the trial was repeated. A "choice" was defined as the subject's nose passing over the edge of a bowl. Subjects were required to choose the baited bowl first in four out of five consecutive trials, within a maximum of 20 trials, to advance to the Gesture Comprehension test. The number of trials taken to either reach criteria or max-out in the first non-aborted testing session was recorded as the subject's score for this task - a lower score indicates better performance. However, subjects that failed to meet this criterion within 20 trials in their first session were tested in another session after a break of at least 30 minutes, if time allowed. This allowed for another opportunity to meet criterion and advance to the Gesture Comprehension tests.

Abort Criteria

If the subject did not make a choice within 15 seconds on two trials in a row, or exhibited any signs of significant stress (including excessive whining, barking, escape behavior, or defecation), a break was taken. In most cases where a break was needed, another attempt occurred later the same day. After a break, the subject started over from the beginning of the Phase 1 warm up trials, regardless of how many trials had been completed before the break.

**Gesture Comprehension**

General Procedures

There were four different tasks in this session: arm pointing, novel marker, body vs point, and odor control. Six trials of each task were conducted. The order of the tasks was the same for all subjects. Bowls were placed at both positions R and L, and E was provided a list denoting which side to place the food on for each trial. Side placement was counterbalanced with the constraint that the reward could not be on the same side in more than two consecutive trials. H first centered the subject at the start line (S) and gently held the subject in place. E approached the subject to present the food reward in her hand and said "look!", allowing the puppy to see and sniff the food briefly. E then closed her fingers around the food and rotated her wrist so that the back of her hand faced the subject, occluding the food. With the reward occluded, E then walked backwards to the bowl at position R, bent down and either placed or pretended to place (as appropriate for the trial) the food in the bowl, then walked across to and did the same at the bowl at position L, making identical hand movements and sounds at each bowl. E then knelt behind the center position (Location E) and performed the designated gesture for the task. She then said "okay!", signaling H to let go of the subject. Throughout the trial, H (who did not know the location of the hidden food) sat still in place.

A choice was defined as the subject touching the bowl or the subject's nose passing over the edge of the bowl. If the subject correctly chose the baited bowl, the subject was allowed to eat the food and E gave verbal praise. If the subject chose the incorrect bowl, E said "wrong" in a monotone voice and the subject was not rewarded nor allowed to see where the food was located. If the subject did not make a choice within 15s, the trial was repeated.

*Arm Pointing*

After baiting, E knelt at location E, bent forward to eye level with the subject, and pointed with her proximal arm to the baited bowl, index finger extended and head turned toward the baited bowl. E then turned her head to look at the subject, said "look!", and turned her head back towards the baited bowl gaze alternating this way three times, all while maintaining the pointing arm's position. E then said "Okay!" and maintained the pointing gesture and gazed towards the baited bowl until the trial ended.

*Novel Marker*

After baiting, E knelt at location E, picked up a small blue wooden block (5cm x 5 cm x 5cm) in her right hand, and held it out in front of her to show the subject while saying "look!". From the kneeling position, E reached over to set the block next to the bowl at position R, and either left it there if that bowl was the baited one in the trial, or picked it back up. She then reached over to either touch her empty hand to the spot next to position L if she had left the block at position R, or set the block next to the bowl at position L. This procedure ensured that both bowls had equal attention from the experimenter and proximity to her hand. E then returned her hands to rest on her thighs, said "Okay!", and remained kneeling and looking straight down until the trial ended.

*Body vs Point Control*

This gesture was identical to that in the arm pointing task, except that E1 knelt directly behind the *unbaited* bowl (behind location R or L) instead of in the center (location E). This created a choice between the bowl which E was closest to in proximity, and the bowl towards which E was pointing. This controlled for the possibility that if wolves performed poorly on the arm pointing task, it could be explained simply by avoidance of the human. If wolves were actively avoiding the human, they would choose the baited bowl on this task.

*Odor Control*

No gesture was administered - after baiting, E knelt down in the center (location E) resting her hands on her thighs, said “Okay!”, and remained kneeling and looking straight down until the trial ended. This created a situation in which subjects had no social information to use to find the food, testing for the possibility that they could use solely olfactory information to locate the food.

Abort Criteria

If the subject did not make a choice within 15s on two trials in a row, or exhibited any signs of significant stress including whining, escape behavior, or excessive defecation, a break was taken. In most cases, especially in the middle of a test trial block, another attempt occurred on the same day. In this case, two Working Memory trials were conducted as warm-ups (one on each side). The subject was required to successfully find the reward on both sides to resume test trials. In the case where the session resumed on a subsequent day, another complete Working Memory session was conducted as warm-ups, and subjects were required to reach criterion prior to resuming testing.  In some cases, due to time constraints (i.e. field seasons being over), another attempt was not possible.

### Inhibitory Control

Apparatus

A cylinder was constructed by taping clear flexible plastic around two solid plastic rings (1ft wide x 11in diameter).  A tube of black opaque stretchy fabric was made to fit as a cover. The cylinder and fabric cover both had a strip of Velcro at the bottom, which could attach the cylinder to a strip of Velcro on a square wooden base (2ft x 2ft) to keep the apparatus still if bumped by the subject.

Warm-up Trials - Opaque Cylinder

H centered the subject at the near start line (S’) and gently held the subject in place. E knelt at location E, behind the cylinder which had the opaque cover on and was positioned at location C. E reached forward (over the cylinder) to present the food reward in her right hand and said “look!”, allowing the subject to see and sniff the food briefly.  E then placed the food inside the cylinder, entering it from the right side (the open side facing position R). E then placed her hands on her thighs, looked down, and said “Okay!”, signaling H to let go of the subject. The subject was permitted 15 seconds to retrieve the reward.  E recorded whether the subject first touched or bumped the front of the cylinder in an attempt to retrieve the reward, or went directly around to the side opening. If the subject did not retrieve the reward within 15s, the trial was repeated. Subjects were required to correctly retrieve the reward, by going directly to the side opening without touching or bumping the front of the cylinder, in 4 out of a window of 5 trials before advancing to test trials.

Test Trials - Transparent Cylinder

The test procedure was identical to the warm-up trials, except that the fabric cover was removed so that the cylinder was transparent. As in warm-up trials, E recorded whether the subject first touched or bumped into the front of the cylinder in an attempt to get to the visible reward (incorrect), or went directly around to the side opening to retrieve the reward without bumping into the cylinder (correct). The subject was allowed to retrieve the reward on all trials regardless of the accuracy of their first attempt. Ten trials were conducted.

Abort Criteria

The session was aborted if the subject exhibited significant signs of stress (including excessive whining, barking, escape behavior, or defecation), did not meet the criterion within 25 warm up trials to advance to the test trials, or did not retrieve the reward within 15s on a total of 10 warm-up trials.

**Human Eye Contact**

Apparatus

A clear container with a clear lid that snapped closed (Rubbermaid Brilliance 1.3 cup container or similar) was super glued to a wooden base (2ft x 2ft) so that it could not be picked up by the subject.

Warm-up Trials - Solvable

H centered the subject at the near start line (S') and gently held the subject in place. E knelt at location E, behind the open clear container with the lid off, which was positioned at location C. E reached forward (over the container ) to present the food reward and said "look!", allowing the subject to see and sniff the food briefly. E then placed the food inside the container, and positioned the lid loosely on top such that it could easily be knocked off . E then placed her hands on her thighs, looked down, and said "Okay!", signaling H to let go of the subject. The subject was allowed to knock off the lid and retrieve the reward, at which point E provided verbal praise and the trial ended. If at any point during the trial the subject somehow knocked the lid into a position which made the reward unreachable, E quietly intervened to make the reward accessible again. If the subject did not successfully retrieve the reward within 30s, the trial was repeated. The subject was required to successfully complete 4 warm-up trials before advancing to the test. The first of these started with the lid leaning up against the container, the second with the lid covering about half of the opening of the container, and the third and fourth with the lid covering three-quarters of the opening of the container.

Test Trials - Unsolvable

Test trials were identical to warm up trials except that E sealed the lid onto the container so that it could not be removed by the subject. The subject was given 30 seconds to attempt to access the reward. E and H remained seated in place and continuously looked at the subject (rotating head and body if necessary) during this period. E and H both held silent stopwatches, and used the start/stop buttons in count-up mode to measure the total time that the subject looked at their faces (*11*). After each test trial, E praised the subject, opened the container, and allowed the subject to retrieve the contents. Four test trials were conducted, and the sum of the time spent looking at E and H's faces across the four trials was used as the measure for analysis.

Abort Criteria

The session was aborted if the subject exhibited signs of significant stress (including excessive whining, barking, escape behavior, or defecation), or if the subject did not obtain the reward within 12 attempts during the warm-up trials.

### Coding and Analysis

All trials were videotaped from an angle (or multiple angles) which captured the subject's response as well as the experimenter (E). Subject responses were usually live-coded by E after each trial; in a minority of cases E was unable to live-code (e.g. stopwatch malfunction) and designated the trial to be coded from video. During live coding and upon reviewing the video for temperament and gesture comprehension, responses were seen as unambiguous and reliability coding was deemed unnecessary. All of the wolf and 20% of dog trials in the Human Eye-Contact test, and 20% of all trials in the Inhibitory Control test, were independently coded for reliability from video. The subset of trials coded from video were chosen pseudo-randomly (ensuring that at least one subject from each of the testing populations was included for each test). Reliability was excellent for both the Human Eye-Contact test (Pearson's $r(16) = .92$, $p<.001$) and the Inhibitory Control test (Cohen's $\kappa = .956$, N = 150, $p <.001$).

Statistical analyses were done using R v.3.5.2. For all analyses, species was treated as a categorical variable, and age and trial number were treated as continuous variables. Bowl choice (baited/unbaited) in the gesture comprehension tests, as well as response (touch/no touch) in the temperament and inhibitory control tests, were treated as binary variables. Looking time in the human eye contact test and the number of trials needed to reach criterion in the working memory test were both treated as continuous variables.

A mixed-effects logistic regression model with random intercepts for each subject was used to estimate the effect of species for each condition in the temperament test. This model also accounted for age and interaction type. Welch's unequal variance t-tests were performed to compare species performance on the Working Memory and Human Eye-Contact tests. We used a mixed-effects logistic regression model with random intercepts for each subject to estimate the probabilities of choosing the indicated location for each species in the two Gesture Comprehension tasks and their two controls. To understand how well individual wolves and dogs performed on the gesture comprehension tests, we counted the number of individuals who achieved significant performance at the $p<.05$ level, defined by non-parametric binomial probability test as 10 out of 12 correct. We also used a mixed-effects logistic regression model with random intercepts for each subject to estimate the probabilities of navigating directly to the side opening without first touching the clear cylinder for each species in the Inhibitory Control test.

Binomial logistic regression models, with species as a predictor variable, were performed for both the Gesture Comprehension tests, as well as the Inhibitory Control test, to determine whether species had a significant effect on performance. Linear regression models, with species as a predictor variable, were performed for the working memory test and the human eye contact test, to determine whether species had a significant effect on performance. To determine whether age, sex, trial number, or temperament had a significant effect on performance, these predictor variables were added to the models and models were compared using AIC and $R^2$.

### Results

#### Temperament

Using a linear contrast test on the mixed-effects logistic regression model reveals that relative to a wolf puppy, the dog puppies' odds of touching the unfamiliar human, familiar human, and unfamiliar object, were 30.52 (95% CI 14.48-73.68, $z = -8.374$, $p<.001$), 5.36 (95% CI 2.99-10.06, $z= -5.593$, $p<.001$), and 1.91 (95% CI 1.16-3.20, $z = -2.57$, $p = .01$) times higher respectively. The dog puppies' odds of touching the familiar object were not significantly different than the wolves' (odds ratio = 1.35, 95% CI 0.72-2.57, z = -0.94, p = .34) (**Fig. 1**). The effect of trial number was insignificant using a likelihood ratio test (Chisq= 126.14, df = 3, p < 2.2e-16). The interaction between species and trial type (stationary vs. moving stimulus) was insignificant using a likelihood ratio test (Chisq = 0.4826, df = 1, p = 0.4872). Similarly, the interaction between species and age was insignificant using a likelihood ratio test (Chisq = 1.89, df = 1, p = 0.1692).

Working Memory

A Welch's two sample t-test comparing performance of dog puppies and wolf pups on the working memory task demonstrates that dogs (M = 11.26 trials, SD = 7) and wolves (M = 11.15, SD = 6.4) were not significantly different in the number of trials it took to reach criteria (correct response on 4 out of 5 consecutive trials), t(56.63) = 0.06, p = .95 (**Fig. 2a**).

Gesture Comprehension

a) *Arm Pointing:* The mixed-effects logistic regression model showed that dog puppies were estimated to choose the indicated location 77.99% of the time (95% CI 70.26 – 84.17, z = 6.11) for the pointing gesture. Wolf puppies were estimated to choose the indicated location 62.08% of the time (95% CI 52.86 – 70.51, z = 2.55) for the pointing gesture. The dog puppies as a group performed above chance (50%) for the pointing gesture (p<.001), as did the wolf puppies as a group (p = .011). The dog puppies chose the indicated location significantly more than the wolf puppies did on the arm pointing gesture (p=.006) (**Fig. 2b**). Using a linear contrast test on a binomial logistic regression model reveals that relative to a wolf puppy, a dog puppy's odds of choosing the indicated location for the pointing gesture were 2.08 (95% CI 1.14 – 3.77) times higher (**Fig. S3**).

To account for the possibility of participants learning over the course of the 6 trials, we also compared Trial 1 performance of the two species using a binomial test, and found that 28 out of 31 of dogs (90%) performed correctly on the first trial, which is significantly higher than the null expectation (p < .001). Meanwhile, only 18 out of 26 wolves (69%) performed correctly on the first proximal pointing trial, which is not significantly higher than the null expectation (p = .07). We also modeled the effect of trial number on performance using a binomial logistic regression and found no significant relationship (p = .15) (**Table S7**).

Finally, we used a Welch's two sample t-test to compare the number of "no-choices" (see Methods) that occurred over the course of obtaining the 6 complete arm-pointing trials, and found no significant difference between the dog (M = 0.55, SD = 1.12) and wolf (M = 0.69, SD = 1.44) puppies, t(46.88) = -0.42, p = .68. This indicates that their differing performance on this task cannot be accounted for by differing motivation to participate in the trials.

b) *Arbitrary Marker:* The mixed-effects logistic regression model showed that dog puppies were estimated to choose the indicated location 78.01% of the time (95% CI 70.73 – 83.89, z = 6.46) for the marker gesture. Wolf puppies were estimated to choose the indicated location 57.25% of the time (95% CI 48.59 – 65.50, z = 1.64) for the marker gesture. The dog puppies as a group performed above chance (50%) for the marker gesture (p<.001), but the wolf puppies as a group did not (p = .100). The dog puppies chose the indicated location significantly more than the wolf puppies did on the marker gesture (p <.001) (**Fig. 2b**). Using a linear contrast test on a binomial logistic regression model reveals that relative to a wolf puppy, a dog puppy's odds of choosing the indicated location for the marker gesture were 2.58(95% CI 1.42 – 4.68) times higher (**Fig. S3**).

To account for the possibility of participants learning over the course of the 6 trials, we also compared Trial 1 performance of the two species using a binomial test, and found that 28 out of 31 of dogs (90%) performed correctly on the first trial, which is significantly higher than the null expectation (p < .001). Meanwhile, 15 out of 26 wolves (58%) performed correctly on the first marker cue trial, which is not significantly higher than the null expectation (p = .56). We also modeled the effect of trial number on performance using a binomial logistic regression and found no significant relationship (p = .28) (**Table S8**).

Finally, we used a Welch's two sample t-test to compare the number of "no-choices" (see Methods) that occurred over the course of obtaining the 6 complete arm-pointing trials, and found no significant difference between the dog (M = 0.16, SD = 0.58) and wolf (M = 0.53, SD = 1.14) puppies, t(35.74) = -1.53, p = .13. There is therefore no evidence for differing motivation between the two species as an explanation for their differing performance on this task.

c) *Body vs Point:* The mixed-effects logistic regression model showed that dog puppies were estimated to choose the location indicated by the point 33.01% of the time (95% CI 28.40 – 39.99, z = -2.56). Wolf puppies were estimated to choose the location indicated by the point 45.99% of the time (95% CI 39.25 – 52.87, z = -0.58). Neither dog puppies (p=1.98) nor the wolf puppies (p=1.43) performed significantly different from chance. The performance was also not significantly different between the two species (p = .18) (**Fig. 2b**). Using a linear contrast test on a binomial logistic regression model reveals that relative to a wolf puppy, a dog puppy's odds of choosing the indicated location were not significantly different than a wolf puppy's (odds ratio= 0.67, 95% CI = 0.38 – 1.16, p = .24) (**Fig. S3**).

d) *Odor control:* The mixed-effects logistic regression model showed that dog puppies were estimated to choose the location indicated by the point 51.89% of the time (95% CI 48.21 – 55.55, z = 0.51). Wolf puppies were estimated to choose the location indicated by the point 42.67% of the time (95% CI 38.68 – 46.74, z = -1.78). Neither dog puppies (p=.61) nor the wolf puppies (p=1.92) performed significantly different from chance. The performance was also not significantly different between the two species (p = .09) (**Fig. 2b**). Using a linear contrast test on a binomial logistic regression model reveals that relative to a wolf puppy, a dog puppy's odds of choosing the indicated location were not significantly different than a wolf puppy's (odds ratio= 1.45, 95% CI = 0.84– 2.51, p = .32) (**Fig. S3**).

*Effect of Age on Gesture Comprehension:* We used a Welch's two sample t-test to compare the ages of the participants, and found that there was no significant difference in average age between the dog puppies (M = 10.87 weeks, SD = 3.18) and the wolf pups (M = 11.38 weeks, SD = 1.70), who participated in the social cues test, (t(47.31) = -0.78, p = .44). We considered age as a variable for each of these tests using linear regression model comparison and found that age was not a significant predictor variable and did not improve the models for any of the test outcomes (**Tables S1 and S2**).

*Effect of Temperament on Gesture Comprehension:* We explored the impact of the individual's performance on the temperament tests on their performance on the gesture comprehension tasks using linear regression models. We found that the overall combined score on the temperament tests is a significant predictor of performance on the arm pointing task, with a higher temperament score (i.e. more attraction to objects and humans) slightly increasing the likelihood of a correct response (**Table S9**). However, when added to models which included species as a predictor variable, the temperament score was no longer significant, suggesting that this influence was due to the species difference in temperament rather than capturing the impact of individual variance (**Tables S9, S10**). Temperament scores were not significant predictor variables for the performance on the two control tasks for gesture comprhension.

Inhibitory Control

The mixed-effects logistic regression model showed that dog puppies were estimated to successfully navigate to the side opening without first touching the clear cylinder 74.01% of the time (95% CI 70.71 – 77.62, z = 5.86). Wolf puppies were estimated to do so 66.19% of the time (95% CI 60.59 – 71.37, z = 2.78). The performance was also not significantly different between the two species (p = .19) (**Fig. 2b**). A linear regression model also shows that species was not a significant variable in predicting the outcome of this test (**Table S5**).

Human Eye Contact

A Welch's two sample t-test comparing behavior of dog puppies and wolf pups during an unsolvable task demonstrates that dogs made eye contact with the human experimenter for significantly more time (M = 4.09 seconds, SD = 4.29) than did the wolves (M = 1.47 seconds, SD = 2.18), t(45.90) = 2.83, p = .007 (Fig. 2d). A linear model shows that species is a significant predictor in the number of seconds of eye contact made by the animal (Table S6).

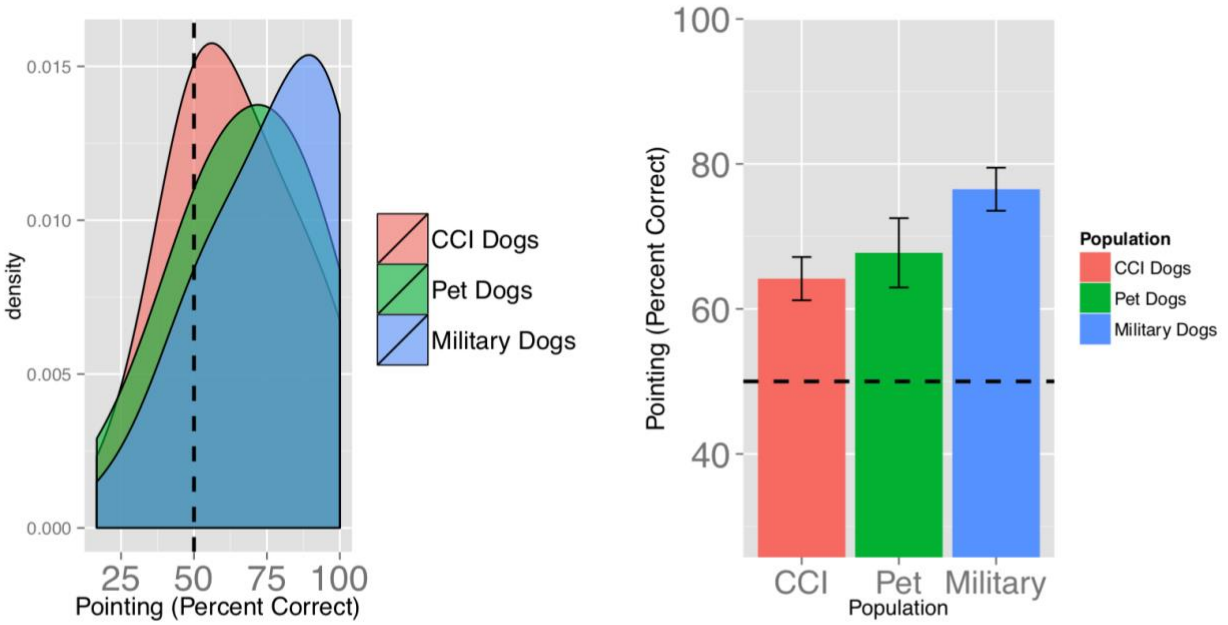


**Fig. S2**. Adult CCI dogs tested for their ability to follow a human pointing gesture in a procedure highly similar to the “proximal pointing” test used here are not more skilled than a heterogeneous sample of pet dogs or military working dogs of similar breed (i.e. Labrador). These CCI dogs were tested around 18-24 months of age just before formal training for working as assistance dogs began (see MacLean & Hare, 2018).

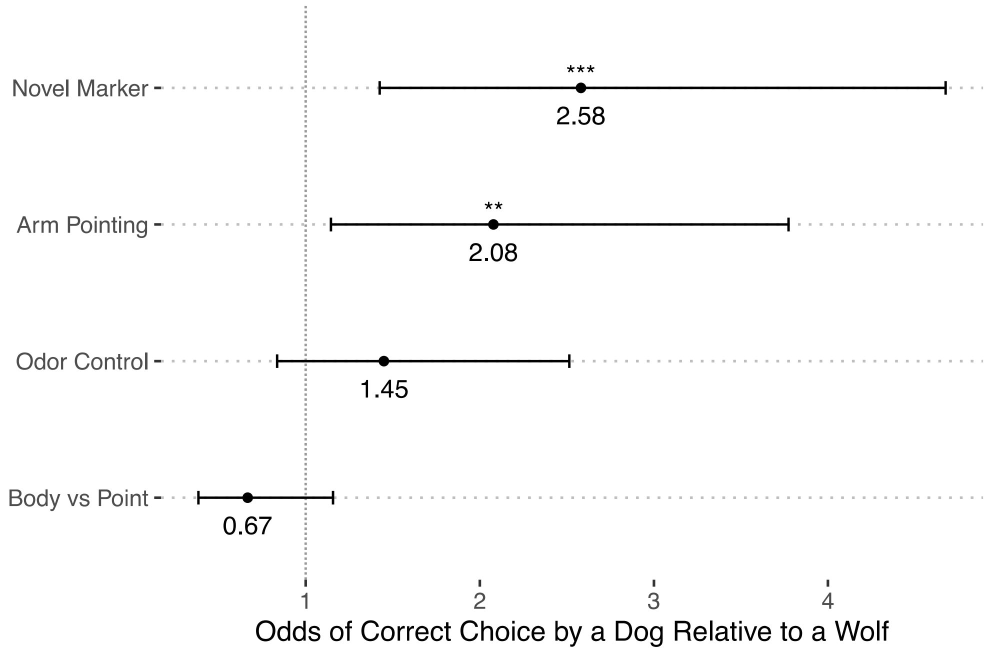


**Fig. S3.** Odds that a dog would choose the baited bowl in each condition as compared to a wolf (i.e., a dog was 2.53 times more likely than a wolf to choose the baited cup in the novel marker task). ** indicates p<.01. Vertical dotted line (odds =1) signifies the point of no difference between species. Bars signify the 95% confidence interval.

<u>Alternative Variable Considerations</u>

*Sex and Age:* We considered sex and age as variables for each of these tests using linear regression model comparison and found that in the tests where a significant species difference was found, species was always the most significant predictor variable for the models (Tables S1, S2, S3, S4, S5, S6).

| | Arm Pointing Models | | |
|---|---|---|---|
| | Species Only | Species + Age | Species + Sex |
| Species = Wolf | -1.082*** | -1.116*** | -1.060*** |
| | (0.302) | (0.303) | (0.308) |
| Age (Weeks) | | 0.067 | |
| | | (0.058) | |
| Sex = F | | | -0.143 |
| | | | (0.308) |
| Constant | 4.774*** | 4.046*** | 4.830*** |
| | (0.204) | (0.667) | (0.238) |
| Observations | 57 | 57 | 57 |
| $R^2$ | 0.189 | 0.208 | 0.192 |
| Adjusted $R^2$ | 0.174 | 0.179 | 0.162 |
| Akaike Inf. Crit. | 180.244 | 180.872 | 182.015 |
| Residual Std. Error | 1.136 (df = 55) | 1.133 (df = 54) | 1.144 (df = 54) |
| F Statistic | 12.829*** (df = 1; 55) | 7.109*** (df = 2; 54) | 6.432*** (df = 2; 54) |
| *Note:* | | | *p<0.1; **p<0.05; ***p<0.01 |

**Table S1.** Linear model comparison for the effect of age and sex on arm pointing gesture comprehension task. For all tables, values presented for each predictor variable are *ß* estimates followed by standard errors in parentheses.

| | Marker Models | | |
|---|---|---|---|
| | Species Only | Species + Age | Species + Sex |
| Species = Wolf | -1.093*** | -1.085*** | -1.004*** |
| | (0.376) | (0.381) | (0.376) |
| Age (Weeks) | | -0.016 | |
| | | (0.074) | |
| Sex = F | | | -0.587 |
| | | | (0.376) |
| Constant | 4.516*** | 4.695*** | 4.744*** |
| | (0.254) | (0.841) | (0.290) |
| Observations | 57 | 57 | 57 |
| $R^2$ | 0.133 | 0.134 | 0.171 |
| Adjusted $R^2$ | 0.117 | 0.102 | 0.140 |
| Akaike Inf. Crit. | 205.278 | 207.226 | 204.754 |
| Residual Std. Error | 1.415 (df = 55) | 1.427 (df = 54) | 1.397 (df = 54) |
| F Statistic | 8.440*** (df = 1; 55) | 4.172** (df = 2; 54) | 5.553*** (df = 2; 54) |
| *Note:* | | | *p<0.1; **p<0.05; ***p<0.01 |

**Table S2.** Linear model comparison for effect of age and sex on marker gesture comprehension task.

| | Total Temperament Models | | |
|---|---|---|---|
| | Species Only | Species + Age | Species + Sex |
| Species = Wolf | -9.318*** | -8.770*** | -9.292*** |
| | (1.067) | (1.063) | (1.079) |
| Age (Weeks) | | -0.449** | |
| | | (0.194) | |
| Sex = F | | | -0.283 |
| | | | (1.050) |
| Constant | 21.711*** | 26.018*** | 21.837*** |
| | (0.661) | (1.972) | (0.813) |
| Observations | 73 | 73 | 73 |
| $R^2$ | 0.518 | 0.552 | 0.518 |
| Adjusted $R^2$ | 0.511 | 0.539 | 0.504 |
| Akaike Inf. Crit. | 428.577 | 425.218 | 430.501 |
| Residual Std. Error | 4.434 (df = 71) | 4.305 (df = 70) | 4.463 (df = 70) |
| F Statistic | 76.228*** (df = 1; 71) | 43.105*** (df = 2; 70) | 37.652*** (df = 2; 70) |
| *Note:* | | | *p<0.1; **p<0.05; ***p<0.01 |

**Table S3.** Linear model comparison for effect of age and sex on total score on all temperament task conditions.

| | Working Memory Models | | |
|---|---|---|---|
| | Species Only | Species + Age | Species + Sex |
| Species = Wolf | -0.102 | 1.584 | -0.379 |
| | (1.685) | (1.564) | (1.666) |
| Age (Weeks) | | $-1.159^{***}$ | |
| | | (0.278) | |
| Sex = F | | | $2.870^{*}$ |
| | | | (1.616) |
| Constant | $11.256^{***}$ | $22.765^{***}$ | $9.988^{***}$ |
| | (1.034) | (2.917) | (1.243) |
| Observations | 69 | 69 | 69 |
| $R^2$ | 0.0001 | 0.208 | 0.046 |
| Adjusted $R^2$ | -0.015 | 0.184 | 0.017 |
| Akaike Inf. Crit. | 463.951 | 462.728 | 462.728 |
| Residual Std. Error | 6.782 (df = 67) | 6.081 (df = 66) | 6.675 (df = 66) |
| F Statistic | 0.004 (df = 1; 67) | $8.664^{***}$ (df = 2; 66) | 1.580 (df = 2; 66) |
| *Note:* | | | $^{*}$p<0.1; $^{**}$p<0.05; $^{***}$p<0.01 |

**Table S4.** Linear model comparison for effect of age and sex on working memory task.

| | Inhibitory Control Models | | |
|---|---|---|---|
| | Species Only | Species + Age | Species + Sex |
| Species = Wolf | -0.670 | -0.759 | -0.664 |
| | (0.581) | (0.556) | (0.585) |
| Age (Weeks) | | $0.222^{**}$ | |
| | | (0.087) | |
| Sex = F | | | 0.179 |
| | | | (0.561) |
| Constant | $7.051^{***}$ | $4.889^{***}$ | $6.969^{***}$ |
| | (0.343) | (0.908) | (0.432) |
| Observations | 60 | 60 | 60 |
| $R^2$ | 0.022 | 0.123 | 0.024 |
| Adjusted $R^2$ | 0.006 | 0.092 | -0.010 |
| Akaike Inf. Crit. | 265.813 | 261.319 | 267.707 |
| Residual Std. Error | 2.145 (df = 58) | 2.050 (df = 57) | 2.162 (df = 57) |
| F Statistic | 1.333 (df = 1; 58) | $3.988^{**}$ (df = 2; 57) | 0.707 (df = 2; 57) |
| *Note:* | | | $^{*}$p<0.1; $^{**}$p<0.05; $^{***}$p<0.01 |

**Table S5.** Linear model comparison for effect of age and sex on inhibitory control task.

| | Human Eye Contact Models | | |
|---|---|---|---|
| | Species Only | Species + Age | Species + Sex |
| Species = Wolf | -2.621** | -4.231*** | -2.684** |
| | (1.173) | (1.294) | (1.171) |
| Age (Weeks) | | 0.413** | |
| | | (0.168) | |
| Sex = F | | | -1.283 |
| | | | (1.119) |
| Constant | 4.091*** | 0.169 | 4.582*** |
| | (0.649) | (1.715) | (0.776) |
| Observations | 49 | 49 | 49 |
| $R^2$ | 0.096 | 0.200 | 0.121 |
| Adjusted $R^2$ | 0.077 | 0.166 | 0.083 |
| Akaike Inf. Crit. | 273.449 | 269.438 | 274.068 |
| Residual Std. Error | 3.785 (df = 47) | 3.598 (df = 46) | 3.772 (df = 46) |
| F Statistic | 4.991** (df = 1; 47) | 5.763*** (df = 2; 46) | 3.170* (df = 2; 46) |
| *Note:* | | | *p<0.1; **p<0.05; ***p<0.01 |

**Table S6.** Linear model comparison for effect of age and sex on human eye contact task.

*Trial Number:* For the Gesture Comprehension tests where dog and wolf performance differed, we considered trial number as a variable using logistic regression model comparison. We found that trial number was not a significant predictor variable, its effect on model fit was minimal when comparing AIC, and while non-significant the coefficient of the association with performance on the gestures tasks is negative (Tables S7, S8).

| | Arm Pointing Models | |
|---|---|---|
| | Species Only | Species + Trial |
| Species = Wolf | -0.732*** | -0.736*** |
| | (0.239) | (0.240) |
| Trial Number | | -0.101 |
| | | (0.070) |
| Constant | 1.202*** | 1.563*** |
| | (0.174) | (0.311) |
| Observations | 342 | 342 |
| Log Likelihood | -204.510 | -203.478 |
| Akaike Inf. Crit. | 413.020 | 412.956 |
| *Note:* | *p<0.1; **p<0.05; ***p<0.01 | |

**Table S7.** Logistic model comparison for effect of trial number on arm pointing gesture comprehension task.

| | Marker Models | |
|---|---|---|
| | Species Only | Species + Trial |
| Species = Wolf | -0.948*** | -0.952*** |
| | (0.239) | (0.239) |
| Trial Number | | -0.075 |
| | | (0.070) |
| Constant | 1.232*** | 1.500*** |
| | (0.175) | (0.308) |
| Observations | 342 | 342 |
| Log Likelihood | -205.928 | -205.345 |
| Akaike Inf. Crit. | 415.856 | 416.690 |
| *Note:* | *p<0.1; **p<0.05; ***p<0.01 | |

**Table S8.** Logistic model comparison for effect of trial number on marker gesture comprehension task.

*Temperament:* For the Gesture Comprehension tests where dog and wolf performance differed, we considered temperament as a variable using logistic regression model comparison. We found that temperament was not a significant predictor for either of the test outcomes when included with species, and adding temperament did not improve the model when comparing AIC (Tables S9, S10).

| | Arm Pointing Models | | |
|---|---|---|---|
| | Species Only | Temperament Only | Species + Temperament |
| Species = Wolf | -1.024*** | | -0.930* |
| | (0.336) | | (0.475) |
| Total Temperament Score | | 0.066** | 0.011 |
| | | (0.029) | (0.040) |
| Constant | 4.774*** | 3.142*** | 4.528*** |
| | (0.210) | (0.572) | (0.900) |
| Observations | 51 | 51 | 51 |
| $R^2$ | 0.160 | 0.094 | 0.161 |
| Adjusted $R^2$ | 0.142 | 0.075 | 0.126 |
| Akaike Inf. Crit. | 164.777 | 168.616 | 166.692 |
| Residual Std. Error | 1.171 (df = 49) | 1.216 (df = 49) | 1.182 (df = 48) |
| F Statistic | 9.303*** (df = 1; 49) | 5.074** (df = 1; 49) | 4.604** (df = 2; 48) |
| *Note:* | | | *p<0.1; **p<0.05; ***p<0.01 |

**Table S9.** Linear model comparison for effect of temperament score on arm pointing gesture comprehension task.

| | Marker Models | | |
|---|---|---|---|
| | Species Only | Temperament Only | Species + Temperament |
| Species = Wolf | -1.016** | | -1.325** |
| | (0.415) | | (0.584) |
| Total Temperament Score | | 0.041 | -0.037 |
| | | (0.037) | (0.049) |
| Constant | 4.516*** | 3.355*** | 5.327*** |
| | (0.260) | (0.713) | (1.107) |
| Observations | 51 | 51 | 51 |
| $R^2$ | 0.109 | 0.025 | 0.119 |
| Adjusted $R^2$ | 0.091 | 0.005 | 0.083 |
| Akaike Inf. Crit. | 186.452 | 191.038 | 187.851 |
| Residual Std. Error | 1.448 (df = 49) | 1.515 (df = 49) | 1.454 (df = 48) |
| F Statistic | 5.986** (df = 1; 49) | 1.257 (df = 1; 49) | 3.251** (df = 2; 48) |
| *Note:* | | | *p<0.1; **p<0.05; ***p<0.01 |

**Table S10.** Linear model comparison for effect of temperament score on marker gesture comprehension task

| *Name* | *Species* | *S ex* | *Pop.* | *Test Year* | *Time with Mother (days)* | *Human Interaction (hr/day)* | *Adult Dog Exposure (hr/day)* | *Testing Room* | *Temp-erament Age (weeks)* | *Working Memory Age (weeks)* | *Gesture Comprehension Age (weeks)* |
|---|---|---|---|---|---|---|---|---|---|---|---|
| *Arya* | Wolf | F | WSC | 2014 | 10-11 | 12 | 12 | Familiar | 13 | 13 | 13 |
| *LittleMan* | Wolf | M | WSC | 2014 | 10-11 | 12 | 12 | Familiar | 13 | 12 | 12 |
| *Shakira* | Wolf | F | WSC | 2014 | 10-11 | 12 | 12 | Familiar | 13 | NA | NA |
| *Honey Pockets* | Wolf | F | WSC | 2014 | 10-11 | 12 | 12 | Familiar | 13 | 12 | 12 |
| *Bran* | Wolf | M | WSC | 2014 | 10-11 | 12 | 12 | Familiar | 7 | 7 | 7 |
| *Sam* | Wolf | M | WSC | 2014 | 10-11 | 12 | 12 | Familiar | 7 | 8 | 8 |
| *Maggie* | Wolf | F | WSC | 2014 | 10-11 | 12 | 12 | Familiar | 13 | 13 | 13 |
| *Graham* | Wolf | M | WSC | 2014 | 10-11 | 12 | 12 | Familiar | 13 | NA | NA |
| *Adam* | Wolf | M | WSC | 2014 | 10-11 | 12 | 12 | Familiar | 7 | NA | NA |
| *Banner* | Wolf | M | WSC | 2016 | 10-11 | 24 | 24 | Familiar | 12 | NA | NA |
| *Skye* | Wolf | F | WSC | 2016 | 10-11 | 24 | 24 | Familiar | 12 | 13 | 13 |
| *Scarlet* | Wolf | F | WSC | 2016 | 10-11 | 24 | 24 | Familiar | 12 | 13 | 13 |
| *Renner* | Wolf | F | WSC | 2016 | 10-11 | 24 | 24 | Familiar | 12 | NA | NA |
| *Alaina* | Wolf | F | WSC | 2016 | 10-11 | 24 | 24 | Familiar | 11 | 12 | 12 |
| *Clay* | Wolf | M | WSC | 2016 | 10-11 | 24 | 24 | Familiar | 11 | 12 | 12 |
| *Cherri* | Wolf | F | WSC | 2016 | 10-11 | 24 | 24 | Familiar | 11 | 12 | 12 |
| *Ezra* | Wolf | F | WSC | 2016 | 10-11 | 24 | 24 | Familiar | 11 | NA | NA |
| *Courtney* | Wolf | F | WSC | 2016 | 10-11 | 24 | 24 | Familiar | 11 | NA | NA |
| *Tiny* | Wolf | F | WSC | 2017 | 10-11 | 24 | 24 | Familiar | 10 | 12 | 12 |
| *Grey* | Wolf | M | WSC | 2017 | 10-11 | 24 | 24 | Familiar | 10 | 12 | 12 |
| *Mara* | Wolf | F | WSC | 2017 | 10-11 | 24 | 24 | Familiar | 8 | 9 | 9 |
| *Dwight* | Wolf | M | WSC | 2017 | 10-11 | 24 | 24 | Familiar | 8 | 9 | 9 |
| *Garland* | Wolf | M | WSC | 2017 | 10-11 | 24 | 24 | Familiar | 11 | 12 | 12 |
| *Kameron* | Wolf | M | WSC | 2017 | 10-11 | 24 | 24 | Familiar | 8 | 9 | 9 |
| *Gavin* | Wolf | M | WSC | 2017 | 10-11 | 24 | 24 | Familiar | 11 | 12 | 12 |
| *Nathan* | Wolf | M | WSC | 2017 | 10-11 | 24 | 24 | Familiar | 11 | 12 | 12 |
| *Jordan* | Wolf | F | WSC | 2017 | 10-11 | 24 | 24 | Familiar | 12 | 13 | 13 |
| *Lexi* | Wolf | F | WSC | 2017 | 10-11 | 24 | 24 | Familiar | 12 | NA | NA |
| *Dwayne* | Wolf | M | WSC | 2018 | 10-11 | 24 | 24 | Familiar | 11 | 12 | 12 |

| | | | | | | | | | | | |
|---|---|---|---|---|---|---|---|---|---|---|---|
| *Artemis* | Wolf | F | WSC | 2018 | 10-11 | 24 | 24 | Familiar | 12 | 12 | 12 |
| *Freya* | Wolf | F | WSC | 2018 | 10-11 | 24 | 24 | Familiar | 12 | 12 | 12 |
| *Jane* | Wolf | F | WSC | 2018 | 10-11 | 24 | 24 | Familiar | 11 | 13 | 13 |
| *Gallo* | Wolf | M | WSC | 2018 | 10-11 | 24 | 24 | Familiar | 10 | 10 | 10 |
| *Madoni* | Wolf | F | WSC | 2018 | 10-11 | 24 | 24 | Familiar | 10 | 10 | 10 |
| *Blue* | Wolf | F | WSC | 2019 | birth | 24 | 24 | Familiar | NA | NA | NA |
| *Echo* | Wolf | M | WSC | 2019 | birth | 24 | 24 | Familiar | NA | NA | NA |
| *Nicholas* | Wolf | M | WSC | 2019 | 10-11 | 24 | 24 | Familiar | NA | NA | NA |
| *Franklin* | Dog | M | P4P | 2016 | 50-60 | 16 | 0 | Familiar | 13 | 13 | 13 |
| *Booth* | Dog | M | P4P | 2016 | 50-60 | 16 | 0 | Familiar | 13 | 13 | 13 |
| *Brooks* | Dog | F | P4P | 2016 | 50-60 | 16 | 0 | Familiar | 12 | 12 | 12 |
| *Hollis* | Dog | F | P4P | 2016 | 50-60 | 16 | 0 | Familiar | 12 | 12 | 12 |
| *Wade* | Dog | F | P4P | 2016 | 50-60 | 16 | 0 | Familiar | 12 | 12 | 12 |
| *Hampshire* | Dog | F | P4P | 2016 | 50-60 | 16 | 0 | Familiar | 11 | 11 | 11 |
| *Penn* | Dog | F | P4P | 2016 | 50-60 | 16 | 0 | Familiar | 11 | 11 | 11 |
| *Porter* | Dog | F | P4P | 2016 | 50-60 | 16 | 0 | Familiar | 11 | 11 | 11 |
| *Divot* | Dog | M | P4P | 2017 | 50-60 | 16 | 0 | Familiar | 13 | 13 | 13 |
| *JGrey* | Dog | M | EENP | 2017 | 50-60 | 16 | 0 | Unfamiliar | 10 | 11 | 11 |
| *Apollo* | Dog | M | EENP | 2017 | 50-60 | 16 | 0 | Unfamiliar | 13 | 13 | 13 |
| *Zelda* | Dog | F | EENP | 2017 | 50-60 | 16 | 0 | Unfamiliar | 13 | 14 | 14 |
| *Orion* | Dog | M | EENP | 2018 | 50-60 | 16 | 0 | Unfamiliar | 16 | 16 | 16 |
| *CCI1* | Dog | M | DPK | 2018 | 50-60 | 16 | <5 | Familiar | 17 | 18 | 18 |
| *CCI2* | Dog | M | DPK | 2018 | 50-60 | 16 | <5 | Familiar | 15 | 15 | 15 |
| *CCI3* | Dog | M | DPK | 2018 | 50-60 | 16 | <5 | Familiar | 13 | 14 | 14 |
| *CCI4* | Dog | M | DPK | 2018 | 50-60 | 16 | <5 | Familiar | 13 | 14 | 14 |
| *CCI5* | Dog | M | DPK | 2018 | 50-60 | 16 | <5 | Familiar | 14 | 15 | 15 |
| *CCI6* | Dog | F | CCI | 2019 | 50-56 | <2 | 24 | Unfamiliar | 7 | 7 | NA |
| *CCI7* | Dog | M | CCI | 2019 | 50-56 | <2 | 24 | Unfamiliar | 7 | 7 | 7 |
| *CCI8* | Dog | M | CCI | 2019 | 50-56 | <2 | 24 | Unfamiliar | 7 | 7 | NA |
| *CCI9* | Dog | F | CCI | 2019 | 50-56 | <2 | 24 | Unfamiliar | 7 | 7 | 7 |
| *CCI10* | Dog | F | CCI | 2019 | 50-56 | <2 | 24 | Unfamiliar | 7 | 7 | 7 |
| *CCI11* | Dog | M | CCI | 2019 | 50-56 | <2 | 24 | Unfamiliar | 7 | 7 | NA |
| *CCI12* | Dog | M | CCI | 2019 | 50-56 | <2 | 24 | Unfamiliar | 7 | 7 | NA |
| *CCI13* | Dog | F | CCI | 2019 | 40-44 | <2 | 0 | Unfamiliar | 7 | 7 | NA |
| *CCI14* | Dog | M | CCI | 2019 | 40-44 | <2 | 0 | Unfamiliar | 7 | 7 | NA |
| *CCI15* | Dog | M | CCI | 2019 | 40-44 | <2 | 0 | Unfamiliar | 7 | 7 | 7 |
| *CCI16* | Dog | F | CCI | 2019 | 40-44 | <2 | 0 | Unfamiliar | 7 | 7 | NA |
| *CCI17* | Dog | F | CCI | 2019 | 40-44 | <2 | 0 | Unfamiliar | 7 | 7 | 7 |
| *CCI18* | Dog | F | CCI | 2019 | 50-56 | <2 | 0 | Unfamiliar | 7 | 8 | NA |
| *CCI19* | Dog | M | CCI | 2019 | 50-56 | <2 | 0 | Unfamiliar | 7 | 8 | 8 |
| *CCI20* | Dog | F | CCI | 2019 | 50-56 | <2 | 0 | Unfamiliar | 7 | 8 | 8 |
| *CCI21* | Dog | M | CCI | 2019 | 50-56 | <2 | 0 | Unfamiliar | 7 | 8 | NA |

| | | | | | | | | | | | |
|---|---|---|---|---|---|---|---|---|---|---|---|
| *CCI22* | Dog | F | CCI | 2019 | 50-60 | <2 | 0 | Unfamiliar | 8 | 8 | NA |
| *CCI23* | Dog | F | CCI | 2019 | 50-60 | <2 | 0 | Unfamiliar | 8 | 8 | NA |
| *CCI24* | Dog | M | CCI | 2019 | 50-60 | <2 | 0 | Unfamiliar | 8 | 8 | 8 |
| *CCI25* | Dog | M | CCI | 2019 | 50-60 | <2 | 0 | Unfamiliar | 8 | 8 | 8 |
| *CCI26* | Dog | M | CCI | 2019 | 50-60 | <2 | 0 | Unfamiliar | 8 | 8 | 8 |
| *CCI27* | Dog | M | CCI | 2019 | 50-60 | <2 | 24 | Unfamiliar | 8 | 8 | 8 |
| *CCI28* | Dog | F | CCI | 2019 | 50-60 | <2 | 24 | Unfamiliar | 8 | 8 | NA |
| *CCI29* | Dog | F | CCI | 2019 | 50-60 | <2 | 24 | Unfamiliar | 8 | 8 | 8 |
| *CCI30* | Dog | F | CCI | 2019 | 50-60 | <2 | 24 | Unfamiliar | 8 | 8 | NA |
| *CCI31* | Dog | M | CCI | 2019 | 50-60 | <2 | 24 | Unfamiliar | 8 | NA | NA |
| *CCI32* | Dog | M | CCI | 2019 | 40-44 | <2 | 0 | Unfamiliar | 8 | 8 | 8 |

**Table S11.** Each subject's name, species, origin (note that DPK puppies were born and raised to weaning at CCI as described in methods), birth year, number of days with mother, daily hours humans were available for physical interaction during rearing, daily hours of exposure to non-maternal adult dogs*, familiarity of testing room, and age in weeks at date of testing for each of the five tests in the battery.
* To the best of our knowledge, based on raiser reports. If an adult dog besides the mother was reported to live in the home of a home-reared dog puppy, we report 24-hour exposure, but we cannot know for sure how often, if at all, the puppy saw or interacted with the adult dog. It is also possible that some home-reared puppies had interactions
